# Daytime light enhances the amplitude of circadian output in a diurnal mammal

**DOI:** 10.1101/2020.06.22.164194

**Authors:** Beatriz Bano-Otalora, Franck Martial, Court Harding, David A. Bechtold, Annette E. Allen, Timothy M. Brown, Mino D. C. Belle, Robert J. Lucas

**Affiliations:** Centre for Biological Timing, Faculty of Biology Medicine & Health, University of Manchester, Manchester, UK; Division of Neuroscience and Experimental Psychology, Faculty of Biology Medicine and Health, University of Manchester, UK; Division of Diabetes, Endocrinology and Gastroenterology, Faculty of Biology Medicine and Health, University of Manchester, UK; Institute of Biomedical and Clinical Sciences, University of Exeter Medical School, University of Exeter, UK

**Keywords:** diurnality, circadian rhythms, daylight, suprachiasmatic nucleus, amplitude, electrical activity

## Abstract

Mammalian circadian rhythms are orchestrated by a master pacemaker in the hypothalamic suprachiasmatic nuclei (SCN), which receives information about the 24 h light:dark cycle from the retina. The accepted function of this light signal is to reset circadian phase in order to ensure appropriate synchronisation with the celestial day. Here, we ask whether light also impacts another key property of the circadian oscillation, its amplitude. To this end, we measured rhythms in behavioural activity and body temperature, and SCN electrophysiological activity in the diurnal murid rodent *Rhabdomys pumilio* following stable entrainment to 12:12 light:dark cycles at 4 different daytime intensities (ranging from 12.77 to 14.80 log melanopsin effective photons/cm^2^/s). *Rhabdomys* showed strongly diurnal activity and body temperature rhythms in all conditions, but measures of rhythm robustness were positively correlated with daytime irradiance under both entrainment and subsequent free run. Whole-cell and extracellular recordings of electrophysiological activity in *ex vivo* SCN revealed substantial differences in electrophysiological activity between dim and bright light conditions. At lower daytime irradiance, daytime peaks in SCN spontaneous firing rate and membrane depolarisation were substantially depressed, leading to an overall marked reduction in the amplitude of circadian rhythms in spontaneous activity. Our data reveal a previously unappreciated impact of daytime light intensity on SCN physiology and the amplitude of circadian rhythms, and highlight the potential importance of daytime light exposure for circadian health.

## Introduction

In mammals, near 24h (circadian) rhythms in physiology and behaviour are orchestrated by a master clock located in the hypothalamic suprachiasmatic nuclei (SCN) (1, 2). The SCN clock generates a circadian rhythm in electrical activity, with neurons significantly more excited during the day (up-state) than at night (down-state) (3, 4). This endogenous rhythm is synchronised (entrained) to the external 24h light-dark cycle via input from the retina (5). Thus, light exposure in the circadian night induces adjustments in circadian phase to ensure that internal time faithfully reflects external (celestial) time. Conceptual and mathematical models of light’s impact on the clock address this ability to reset circadian phase (the basis of entrainment) (6-9). However, there is growing interest in the possibility that another fundamental property of circadian rhythms, their amplitude, may also be influenced by light.

It is well established that higher daytime light exposure can increase the amplitude and reliability of 24hr rhythms in some aspects of physiology and behaviour (10-12). However, such effects have typically been attributed to the ability of light to directly engage some of the systems under circadian control (e.g. increasing alertness and body temperature) (13-15) and thus enhance rhythm amplitude *de facto*, without impacting the circadian clock itself. However, reports that enhanced daytime light can also lead to higher production of melatonin on the subsequent night, many hours after light exposure has ceased (12, 16) pose a challenge to that explanation. The long-lasting nature of that effect raises the possibility that daytime light may impact circadian amplitude in a way that cannot simply be accounted for by the immediate effects of light on physiological outputs. We set out here to address this possibility by asking whether increasing daytime irradiance could produce a persistent alteration in the amplitude of circadian rhythms at the whole animal level and whether this could be traced back to changes in the SCN circadian oscillator itself.

A challenge to studying the impact of daytime light exposure in common laboratory models (mice and rats) is that they are nocturnal and employ strategies to avoid light in the day (such as curling up asleep). We therefore used a diurnal rodent, *Rhabdomys pumilio* (the four striped mouse) (17-19) which is active through the day in both the lab and wild, ensuring good exposure to modulations in daytime light. We find that increasing irradiance across a range equivalent to that from dim indoor lighting to natural daylight enhances the reproducibility and robustness of behavioural and physiological rhythms at the whole animal level. This effect is associated with profound differences in the electrophysiological activity of the SCN, with bright daytime light producing persistent increases in SCN excitability and enhancing the amplitude of the circadian variation in spontaneous neuronal activity.

## Results

### Enhancing daytime irradiance increases reproducibility and robustness of circadian rhythms

We first set out to determine the impact of increasing daytime light intensity on circadian rhythms in behaviour (general locomotor activity and voluntary wheel running activity) and physiology (body temperature, Tb) under stable entrainment to a 12:12 light:dark (LD) cycle. As a previous study failed to identify a pronounced impact on rhythmicity across a range of lower illuminances (19), here we applied daytime irradiances (Fig.1A) extending into the lower portion of the daylight range [12.77 to 14.80 log melanopsin effective photons/cm^2^/s; or Melanopic EDI (equivalent daylight illuminance) of 17.92 to 1941.7 lx]. Lighting conditions aimed to reproduce the *Rhabdomys* experience of natural daylight by approximating the relative activation for melanopsin, rod opsin and cone opsins (Fig.1A; although note that we were unable to recreate levels of near UV required to adequately stimulate S-cones). When exposed to these lighting conditions, all animals remained entrained to the 24h light-dark cycle (Fig.1B).

**Figure 1.**
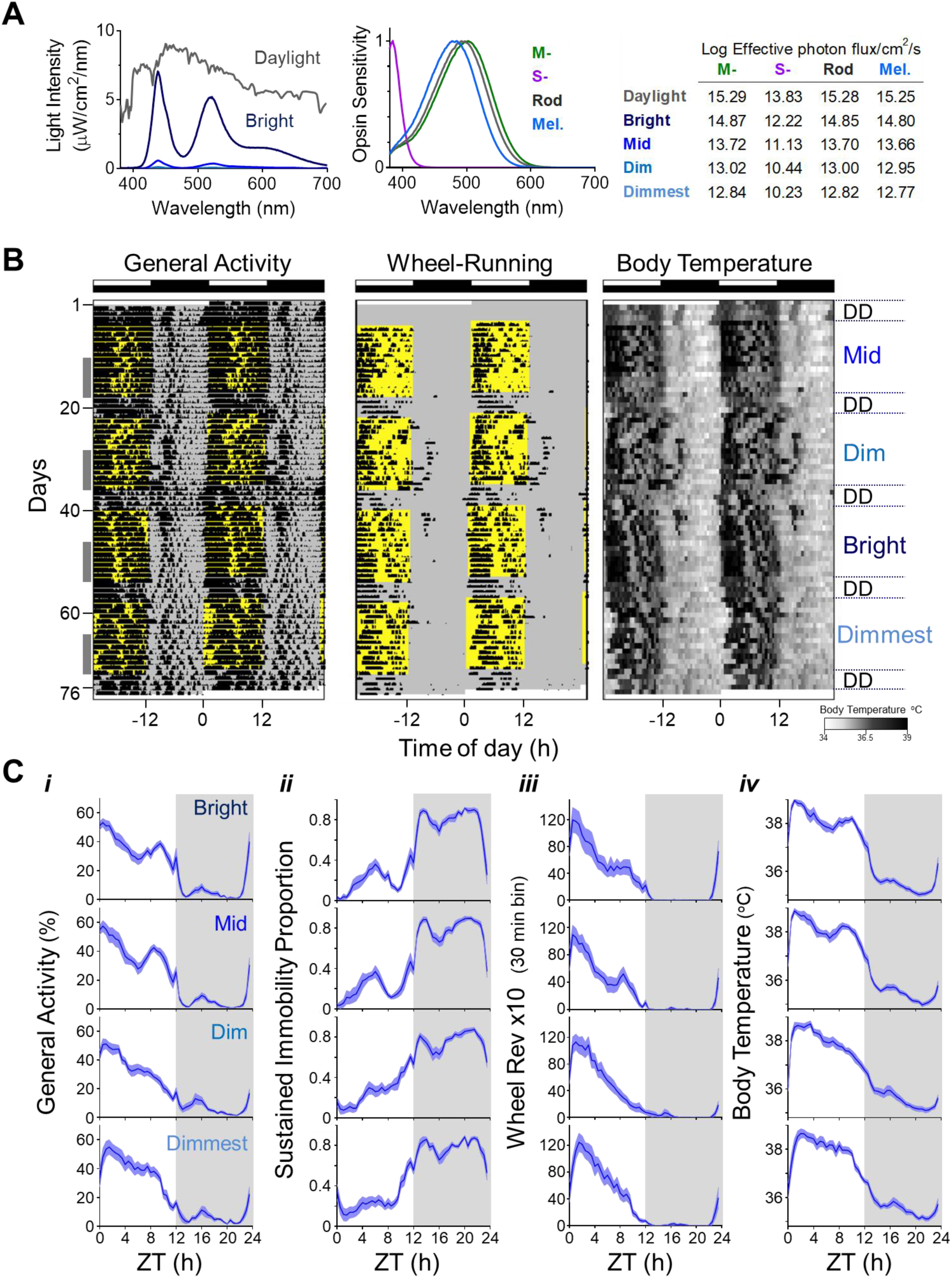
Impact of increasing daytime light intensity on circadian rhythms in the Rhabdomys. **(A)** Lighting conditions. Left panel shows the spectral power distribution of our light source at different irradiances and that of daylight on an overcast day. Middle panel shows expected spectral sensitivity profile of mammalian rod opsin and melanopsin and *Rhabdomys* MWS- & SWS - cone opsins (43) corrected for lens transmission. Right table shows Log_10_ effective photon fluxes for each opsin across the different lighting stimuli used (Bright, Mid, Dim, Dimmest). **(B)** Double-plotted actograms for general activity, wheel running and body temperature (scale below) from a representative *Rhabdomys* under different lighting conditions, over a period of 2.5 months. Time of light exposure is indicated in yellow, and intensity of the light is shown on the left. Entrainment to 12:12 LD cycle at each irradiance ran for 2 weeks followed by 4 days in constant darkness (DD). Note that across conditions, activity is largely restricted to daytime (light phase) coinciding with higher body temperature values. However, the rhythms become less robust at low light levels. Grey columns on the right indicate the 8-day period at the end of each stage used for analysis reported in Supplementary Table1 and Fig.2. **(C)** Mean waveforms for the recorded biological rhythms (***i*:** general activity; ***ii*:** sustained immobility; ***iii*:** voluntary wheel running activity; ***iv*:** body temperature) across the different lighting conditions. Mean waveforms under bright intensities are shown in the top panels and the dimmest conditions in panels at the bottom. Values are expressed as mean ± SEM (n=12), grey areas indicate period of darkness. ZT0: Zeitgeber Time 0 corresponds to time of lights on.

Rhabdomys were strongly diurnal under all light conditions, spending most of the day awake and active, and being quiescent at night (Fig. 1B-C and Supplementary Table 1). Indeed, there was no significant effect of daytime irradiance on the proportion of activity occurring during the light period, nor did daytime irradiance alter mean Tb or the total amount of general or wheel running activity across the 24h (Table 1). However, at a finer timescale, activity patterns were impacted by light intensity. Thus, when we quantified the appearance of periods of sustained immobility (which in mice correspond to episodes of sleep) we found these occurred less often during the day (Supplementary Table 1) and were less fragmented (lower intradaily variability; Fig. 2A) under higher irradiance. Moreover, the day-to-day reproducibility of this rhythm in immobility was also positively correlated with irradiance (Fig.2B). These findings imply that rhythmicity in this proxy for sleep is more robust under higher light. That conclusion was supported by analysis of Tb rhythms, which also showed negative correlations between daytime irradiance and fragmentation and day-to-day variability (Fig.2C&D).

**Figure 2.**
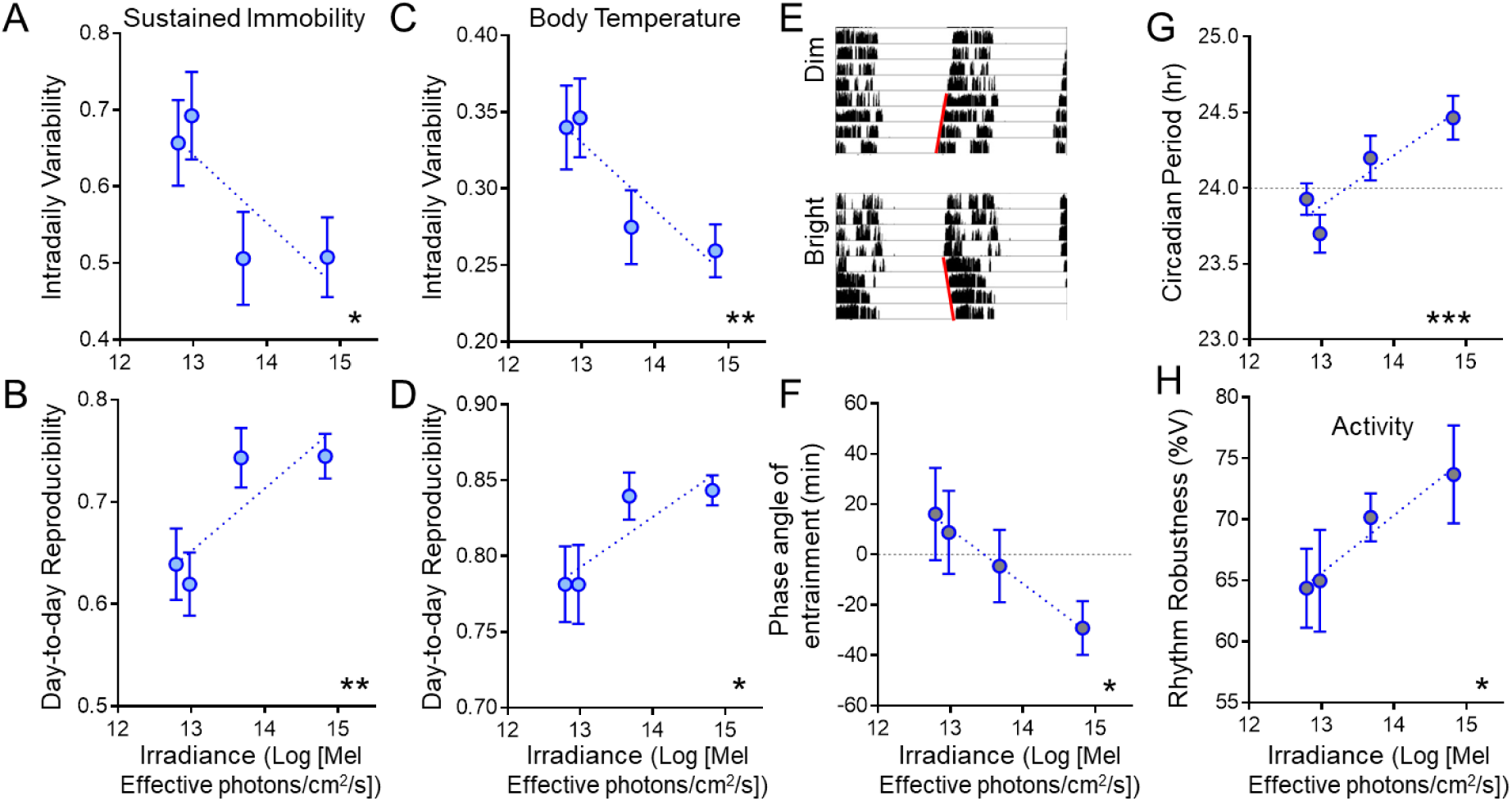
Impact of increasing daytime light intensity on reproducibility and robustness of circadian rhythms under entrained and subsequent free-running conditions. **(A)** Intra-daily variability and **(B)** day-to-day reproducibility of the sustained immobility rhythm as a function of daytime irradiance under entrained conditions. **(C&D)** Same as in A&B but for the body temperature rhythm. **(E)** Representative wheel running activity actogram for a *Rhabdomys* over the last 4 days of entrainment under dim (top) or bright (bottom) irradiance, and subsequent 4 days of free run in constant darkness (note difference in slope of red lines fit to activity onsets in free run between conditions). **(F)** Relationship between phase angle of entrainment for wheel running activity rhythm and daytime irradiance. **(G)** Free-running period and **(H)** robustness of the activity rhythm across all animals as a function of the prior entraining irradiance. Values are expressed as mean ± SEM (n=12, A-D; n=11, F-H); significant relationships identified by linear regression (*p < 0.05, **p < 0.01, and ***p < 0.001).

As rhythms under entrained conditions may reflect direct effects of the LD cycle as well as the output of the circadian clock, we analysed rhythms expressed over 4 days of constant darkness at the end of each light condition to reveal the characteristics of the clock in the absence of direct environmental drive. In all conditions, activity rhythms free-ran from the phase adopted during the prior LD cycle, confirming that they had indeed been photoentrained (Fig.2E). Circadian formalisms predict that increasing the amplitude of an entraining stimulus can alter the phase of entrainment (7), and indeed that proved to be the case in these experiments, with rhythms becoming delayed with respect to the prior light:dark cycle at lower intensities (Fig.2F). Analysis of these free-running rhythms revealed effects of daytime light intensity on further circadian properties. Firstly, free-running period was positively correlated with prior irradiance (Fig.2G). Secondly, the %variance in activity explained by the circadian rhythm (an indication of rhythm robustness) was higher in animals previously held under brighter conditions (Fig.2H).

### Impact of daytime irradiance on SCN electrical activity

The effects of prior light exposure on rhythms in subsequent constant darkness indicate a persistent effect of daytime light intensity on circadian physiology. To explore this possibility we turned to an assessment of the central SCN circadian clock itself. To this end, we collected the SCN of animals housed in either bright (n=4) or dim (n=4) daytime irradiances for *ex vivo* electrophysiological recordings.

We first recorded extracellular activity in the SCN region using multi-electrode arrays (MEAs) (Fig.3A). This method allows long-term recordings (24 h) to reveal rhythms in spontaneous firing activity. We observed rhythms in spike firing at electrode sites falling within the SCN and, on the average, spontaneous firing rates were higher during the day than at night (Fig.3B). However, there was a dramatic difference in the activity of SCNs from dim vs bright conditions (Fig.3C-G). Firstly, overall spontaneous firing activity was strongly influenced by prior daytime irradiance, with mean multiunit activity across the 24h recording epoch significantly enhanced following the bright LD cycle (Fig. 3C, p<0.001, Mann-Whitney U test). Secondly, rhythmicity was markedly less pronounced in the dim condition. Thus, the fraction of MEA recording sites within the SCN at which a circadian variation in multiunit activity was detected was significantly reduced under the dim condition (Bright: n=58/92,∼63% vs. Dim: n=42/99, ∼42%; 4 slices per condition; p<0.01 Fisher’s Exact Test), Fig. 3D). Moreover, while there was a clear daytime peak in spontaneous activity under both conditions (Fig. 3E&F), the amplitude of that rhythm (difference in firing between peak and trough) was substantially reduced in SCNs taken from animals previously held under dim conditions (Fig.3G; p<0.001, Mann-Whitney U test).

**Figure 3.**
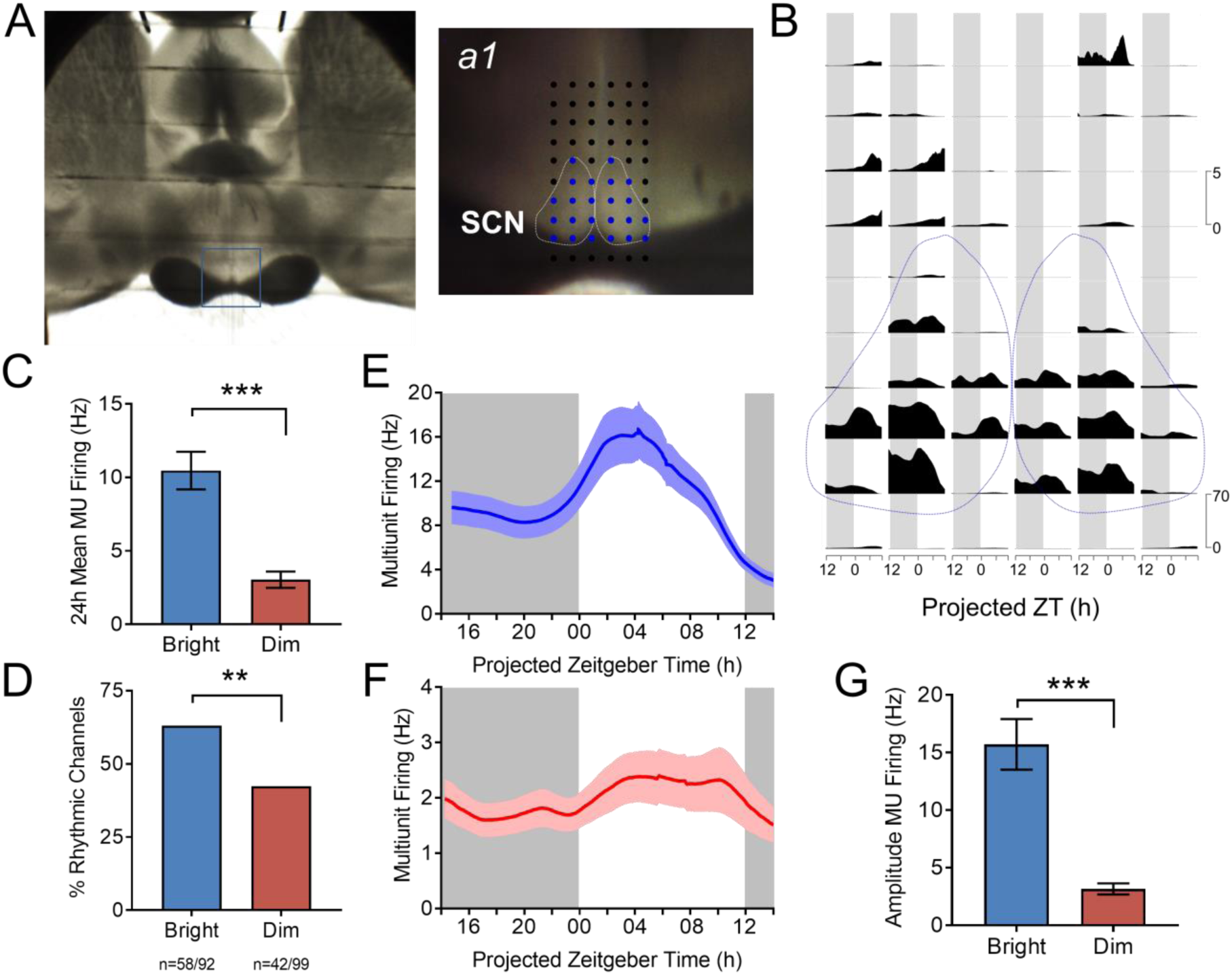
Impact of daytime irradiance on SCN activity. **(A)** Image of a brain slice of *Rhabdomys* placed over the electrode terminals of a 60 channel array to record neuronal electrical activity in the SCN. The recording site is shown in close up to right (a1), with dotted white lines delineating the SCN area and electrodes designated as within the SCN shown in blue. **(B)** Multiunit neuronal activity (MUA) recorded at the array of electrodes for the preparation shown in **A** as a function of projected Zeitgeber time (ZT) for a 24h recording epoch. Spontaneous activity within the SCN (bounded by dotted lines) is higher than surrounding hypothalamus and plotted on a different scale (scale bars to right) **(C)** Time averaged firing rate (MUA) across 24h recording epochs from all channels covering SCN harvested from bright (n=92 channels from 4 slices) vs dim (n=99 channels from 4 slices) conditions. **(D)** Percentage of channels within the SCN classified as rhythmic for each experimental condition (bright vs dim, **p<0.01; Fisher’s exact test, data from 4 slices per condition). **(E&F)** Multiunit firing rate as a function of projected Zeitgeber time for rhythmic recording sites within the SCN from animals maintained under bright (**E**) and dim (**F**) daytime conditions (Bright, n=58; dim, n=42 channels from a total of 4 animals per condition). **(G)** Peak-trough amplitude in the multiunit firing rhythm was significantly lower under dimmer lights. (***p<0.001, Mann-Whitney U-test, bright, n=58; dim, n=42 channels). Grey background indicates the projected period of lights off (ZT12-ZT24). Data are expressed as mean ± SEM.

### Impact of daytime irradiance on single cell neurophysiology in the SCN

The MEA recordings thus reveal that the brighter daytime light exposure produces an SCN that is overall more active and more highly rhythmic, and suggest that daytime irradiance defines important aspects of the SCN network. To explore the nature of this effect, we applied whole-cell patch recordings across the day and at night in order to determine how changes in daytime irradiance impact SCN neurophysiology at the single cell level (Fig.4A-B).

Across the sample of neurons recorded from both bright and dim conditions (n=111 and 104, respectively, 8 Rhabdomys per condition), we found that the resting membrane potential (RMP) ranged from −62.5mV to −21.6mV. Across most of this range, the RMP correlated with firing rate, with more hyperpolarized neurons showing lower spontaneous activity, and at the extreme being silent (Fig.4C-D). Although depolarization generally correlated with higher firing, this was not true for the most depolarized neurons. Such cells showed depolarized low-amplitude membrane oscillations (DLAMOs) in place of spikes or, at the extreme, were depolarized-silent (‘hyper-excited’ (20, 21) (Fig. 4C). Neurons in each of these neurophysiological states were recorded in both conditions; however, their distribution was different. In the bright day group, hyperpolarized silent neurons only appeared at night, with the daytime state being characterised by either firing or highly depolarized cells (Fig. 4E). By contrast, in the dim light exposed group, a subset of neurons were hyperpolarized silent even during the day (Fig. 4E, Day: Bright vs Dim, χ^2^=9.322, p=0.009).

**Figure 4.**
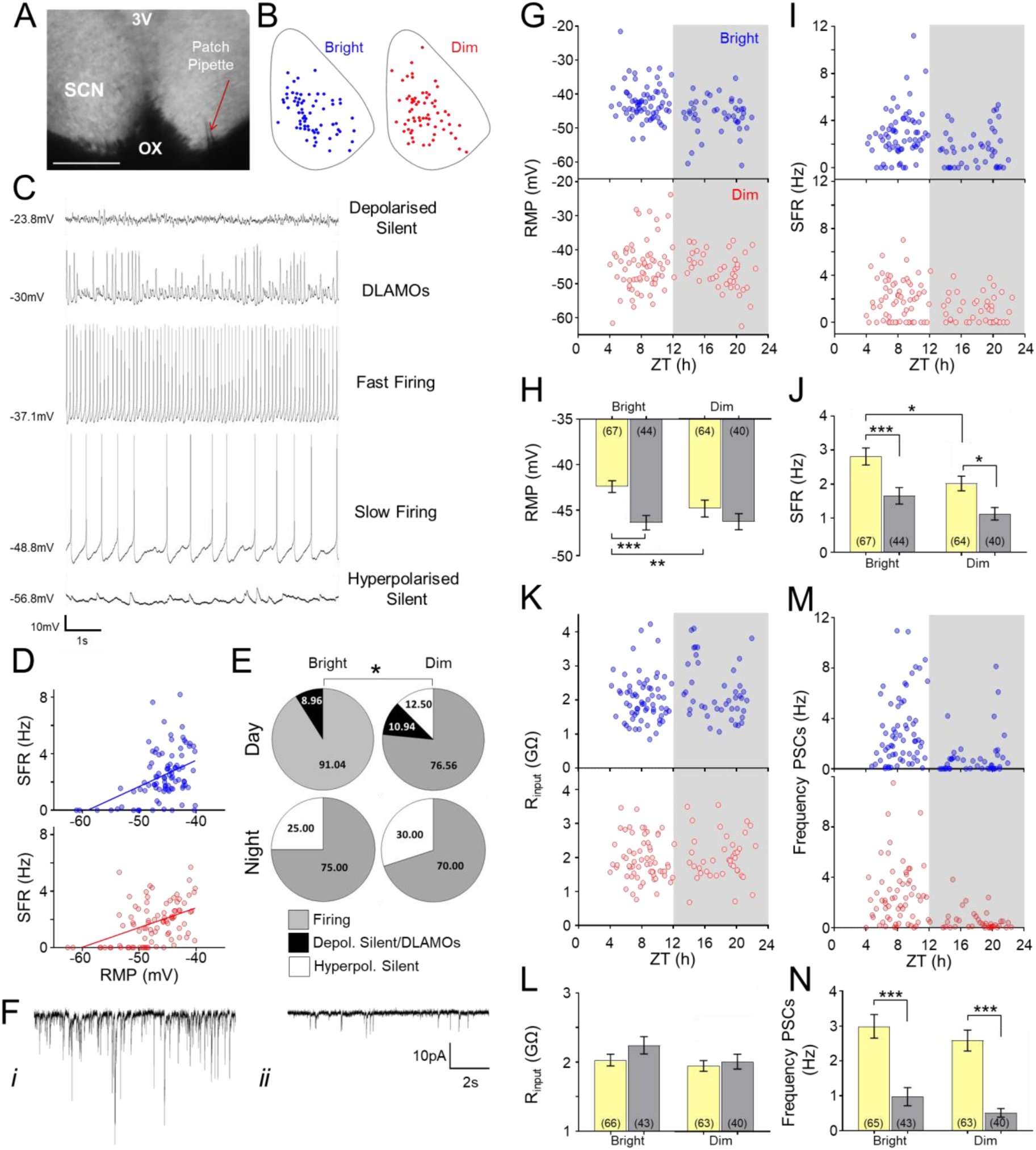
Increasing daytime light intensity impacts SCN neurophysiology at the single cell level. **(A)** Whole-cell recording setup showing bright-field image (4X) of a living SCN in coronal brain slice either side of the third ventricle (3V) and above the optic chiasm (OX), with a patch pipette (red arrow) targeting a SCN neuron. Scale bar 250µm **(B)** Schematic diagram showing the approximate location within a representative SCN coronal slice of the recorded cells giving rise to the daytime datasets under bright (red) or dim (blue) daytime irradiance. Note the similar spatial distribution between experimental groups. **(C)** Representative traces for each of the spontaneous membrane excitability states encountered in *Rhabdomys* SCN neurons (from top): highly depolarized silent; depolarized displaying low-amplitude membrane oscillations (DLAMOs); moderate resting membrane potential (RMP) and firing at high or low rate; and, hyperpolarized silent. **(D)** Positive correlation between RMP and spontaneous firing rate (SFR) for cells recorded from bright (blue) and dim (red) conditions. Cells resting at RMP > −40mV were excluded. p<0.0001, linear regression analysis. **(E)** Pie charts showing the percentages of cells in the different electrophysiological states during the day and at night from bright and dim conditions. (*p=0.01, Chi-Squared test). **(F)** Example of traces from two SCN neurons (voltage-clamped at −70mV) showing post-synaptic currents (PSCs) recorded during the day (***i***) and at night (***ii***). (***i***) Cell displaying high frequency and amplitude PSCs; (***ii***) cell showing low frequency and amplitude PSCs. Scatter plot showing the RMP **(G)**, SFR **(I)**, input resistance (Rinput) **(K)**, and synaptic activity frequency **(M)** of the different cells across zeitgeber times (ZT) for bright (red) and dim (blue) conditions. Each dot represents an individual cell. Grey background indicates the projected phase of lights off. Mean RMP **(H)**, SFR **(J)**, Rinput **(L)**, and synaptic activity frequency **(N)** during the day and at night for SCN neurons from bright or dim conditions. Data in bar charts are expressed as mean ± SEM. Number of cells used are indicated between brackets in each bar. *p < 0.05, **p < 0.01, and ***p < 0.001, Mann-Whitney U-test.

Leaving aside these categorisations, both RMP and spontaneous firing rate (SFR) showed a strong circadian variation in the bright daytime group (Fig. 4G-J). Consistent with our multiunit data and previous reports from other species (22-26), SFR was higher during the day in these animals (p=0.001, Mann-Whitney U). By contrast, rhythms in SFR and RMP were substantially dampened in the dim day group (Fig4. G-J) to the extent that the variation in RMP no longer reached statistical significance. Overall, SFR and RMP were higher in the bright daytime group, especially during the day phase (Fig4.H&J, p= 0.026 and p=0.009 for SFR and RMP, respectively, Mann-Whitney U).

Differences in spontaneous activity and RMP between conditions could originate with disparities in intrinsic properties of the neurones and/or in the degree of extrinsic synaptic input. To evaluate these possibilities, we used voltage clamp mode to record synaptic activity (Fig.4F, n= 108 and 103 from dim and bright daytime conditions, respectively). Overall, we found higher frequency and amplitude of post synaptic currents (PSCs) during the day than at night (p<0.001, Mann-Whitney U). However, both frequency and amplitude of synaptic inputs were similar between conditions (Fig.4M-N), turning the focus to alterations in intrinsic neuronal properties as a likely cause of the difference in spontaneous activity. We therefore turned to measure an overt intrinsic determinant of neuronal excitability – input resistance (Rinput). We found no differences in this parameter between experimental groups either (Fig. 4K-L). However, we did find that Rinput was high in Rhabdomys SCN (as previously reported for other species (20)), highlighting the potential for small changes in intrinsic currents to produce large modulations in RMP.

## Discussion

Our experiments with *Rhabdomys* reveal a fundamental and previously unappreciated effect of daytime irradiance on the SCN circadian clock. Higher daytime light intensity increased the amplitude of the SCN’s daily peak in SFR and neuronal depolarization, to enhance the amplitude of its circadian rhythm in spontaneous activity. This was associated with improvements in measures of rhythm robustness at whole animal level. These effects reflected a genuine long-lasting change in circadian physiology, not simply a direct and immediate response to brighter light, as they persisted once the light stimulus was removed. The effect of daytime light intensity on rhythm amplitude and robustness reported here is not predicted by current theoretical frameworks or mathematical models of light’s impact on the clock. Light is principally considered a modulator of circadian phase (6, 8, 9). In this capacity, light may impact amplitude transiently following disruptive changes in phase or, under very abnormal exposure patterns, by driving the oscillator towards a point of singularity (27). Neither of those effects should be engaged under the conditions of stable entrainment employed here. Under stable entrainment the phase shifting effects of light can result in a correlation between daytime light intensity and the phase relationship between the endogenous clock and the entraining light cycle (termed the phase angle of entrainment). If the period of the entraining light cycle is substantially divergent from 24hr, phase angles of entrainment can be large (and themselves induce pathology (28)). However, phase angles of entrainment should be less extreme when entraining and endogenous periods are fairly similar, and, indeed, here we found only a modest change in phase angle (<1hr) across a wide range of daytime irradiances. The marked relationship between daytime irradiance and circadian amplitude we observe thus implies an additional impact of light exposure on the circadian machinery, distinct from its phase resetting function.

The most striking effect of changing entraining light intensity is on the SCN’s daytime peak in spontaneous electrical activity (‘up state’). Multiunit recordings reveal a dramatic reduction in daytime spiking activity under entrainment to dim daytime light. This effect is also apparent in the whole-cell recordings, in which it is revealed to originate with a shift to more hyperpolarized resting membrane potentials, and the anomalous appearance of electrically silent neurons during the day phase. It is less clear whether entraining irradiance also impacts SCN activity during its nocturnal ‘down state’. MEA recordings revealed a significant reduction in MUA over the subjective night, but this was not apparent in the whole-cell recordings. As many more neurons likely contribute to the MUA recordings it seems likely that there is a modest decrease also in night time activity that falls below our ability to detect in whole-cell recordings. Indeed, such a difference is a predicted consequence of reduced amplitude in the circadian peak in spontaneous activity, as peaks in firing of individual SCN neurons do extend into the circadian night (26). Nevertheless, because the impact on daytime neuronal activity is so much larger, the overall effect is a substantially blunted circadian amplitude. Thus, both MEA and whole-cell recordings reveal that circadian differences in SFR and RMP at the population level are significantly smaller in the dim condition.

As experimental alterations in SCN electrical activity can influence the amplitude of circadian rhythms (29, 30), a parsimonious explanation for our findings is that the reduction in daytime depolarization is a primary impact of changing LD cycle amplitude, with knock-on impacts on the amplitude of SCN firing rate rhythms and the robustness of rhythms at the whole animal level. Future work will be required to determine how the difference in daytime RMP arises. Our whole-cell recordings preclude overall changes in input resistance or the frequency/amplitude of synaptic input as origins for this effect. This suggests a change in the magnitude of some other subtle depolarising inward cation currents acting as a driving force to promote excitability (31). Indeed, the high input resistance of SCN neurones (20) (including those of *Rhabdomys*) means that even small differences in a depolarizing current could result in physiologically relevant changes in RMP. Another possibility, and one which cannot be determined with our methodology, is that exposure to different light intensities may alter the polarity of GABAergic activity in the SCN. GABA is the main neurotransmitter in the SCN, and the balance between its excitatory and inhibitory effects is key in shaping excitability and neuronal network dynamics in this brain region (32). Indeed, shifts in the inhibitory/excitatory ratio of GABAergic activity is well known to occurred with environmental changes such as photoperiods, where increased light exposure favours the excitatory actions of GABA (33).

Importantly, the difference in daytime RMP and SFR is not simply an acute response to the alteration in light exposure. The primary neurotransmitter of the retinohypothalamic tract is glutamate (34), but the SCN network principally employs GABA. Accordingly, in other species, individual SCN neurons can respond to light exposure either with inhibition or excitation (35). Although the literature is sparse, inhibitory responses may be more common in diurnal than nocturnal species (36, 37). Nevertheless, even if we accept that the immediate effect of light is mostly to excite the Rhabdomys SCN, that cannot explain our observations as both our MEA and whole-cell recordings are undertaken at least several hours after any direct drive from the retina has been removed (38). This implies that the effect of changing daytime irradiance on SCN neurophysiology is long lasting. That conclusion is supported by our finding that effects of the prior LD cycle on activity rhythms persist into subsequent free-run in constant darkness.

The question of how brighter daytime light impacts the circadian clock is of more than academic interest. The potential for light exposure at night to disrupt circadian clocks (with potentially wide-ranging impacts on physiology and well-being) is well established (39). The demonstration here that lower daytime irradiance can result in reduced circadian amplitude provides new motivation to consider also the importance of daytime light exposure. The light intensities employed here span a range of plausible indoor light intensities. The lowest (corresponding to around 18 lx photopic illuminance of natural daylight or 40 lx photopic illuminance for a typical 3000K fluorescent source) would be that of a dimly lit room, while the brightest (equivalent of around 1760.8 lx) of a bright room, likely with substantial natural light. Humans increasingly spend much of their day indoors exposed to light intensities far below those typical of outdoor sunlight. This is particularly the case for some of the most vulnerable population groups, including hospital inpatients and care home residents (who may have a predisposition for low circadian amplitude (40-42)). Our findings suggest that, by reducing circadian amplitude, dim daytime light could contribute to some of the disruptions in sleep and circadian organisation that such people suffer. As there is evidence that low amplitude also makes circadian clocks more susceptible to the disruptive effects of nocturnal light, it could also contribute to the widespread occurrence of social jetlag and chronic sleep deprivation in the general population. More optimistically, our data provide a new mechanistic justification for using bright daytime light exposure for therapeutic purposes.

## Methods

### Animals

All animal use was in accordance with the UK Animals, Scientific Procedures Act of 1986, and was approved by the University of Manchester Ethics committee.

A total of 36 adults Rhabdomys (male and female, age 3-9 months) were used. Unless otherwise stated, animals were individually housed under a 12:12h light dark cycle and 22°C ambient temperature in light tight cabinets. Zeitgeber Time (ZT) 0 corresponds to the time of lights on, and ZT12 to lights off. Food and water were available *ad libitum*. Cages were equipped with running wheels (8.8 cm radius) for environmental enrichment.

### Lighting conditions

Light was measured using a calibrated spectroradiometer (SpectroCAL MSII, Cambridge Research Systems, UK) at the cage level between 380-700nm. Light was provided by smart RGBW bulbs (LIFX A60; LIFX, Cremorne, Australia) in which LED intensities could be independently controlled. Light stimuli aimed to approximate the relative activation for melanopsin, rods and cones photoreceptors produced by natural daylight in the Rhabdomys. Light intensity was reduced over 2 decades (bright, mid, dim and dimmest) by introducing neutral density filters (Lee Filters) instead of reducing bulbs outputs in order avoid changes in ambient temperature. The effective photon flux for each photoreceptor (Fig.1A) was calculated by multiplying the spectral power distribution of each light stimulus by the spectral sensitivity of each photopigment corrected for the Rhabdomys spectral lens transmission (43). Corresponding Melanopic equivalent daylight illuminance (EDI) lux were calculated using the CIE S 026 Toolbox (44).

### Experimental design for behavioural studies

Animals were housed under a stable 12:12 light dark cycle with daytime light intensities ranging from Bright, 14.80; mid, 13.66; dim, 12.95; to dimmest, 12.77 Log Effective photon flux/cm^2^/s for melanopsin. Each irradiance ran for 2 weeks and ended with 4 days under constant darkness. We studied a total of 12 animals split over 3 different batches in which different daytime irradiances were presented in a pseudorandom order. (Batch 1 (n=5): Mid, Dim, Bright, Dimmest; Batch 2 (n=2): Dimmest, Mid, Bright, Dim; Batch 3 (n=5): Bright, Dim, Mid, Dimmest).

Activity and body temperature rhythms were recorded throughout the whole experiment. Home cage activity was monitored in 10-sec bins using a passive infrared motion sensor (PIR) system, as previously described (45, 46). Wheel running activity was continuously recorded as wheel revolutions/10-sec intervals using a custom-made data acquisition system. Body temperature was recorded every 30-min using a data logger (Maxim, DS1922L-F5; ThermoChron; Data loggers iButton; Maxim Integrated Products, Sunnyvale, CA, USA). iButtons were de-housed and potted with wax (20% Poly(ethylene-co-vinyl acetate) and 80% paraffin mixture) before surgically implantation into the peritoneal cavity (47).

### Brain slice preparation for electrophysiological recordings

Rhabdomys, housed either under bright (n=12) or dim (n=12) daytime conditions for 2-3 weeks, were sedated with isoflurane (Abbott Laboratories) and immediately culled by cervical dislocation during the light phase (beginning of the day or late day). Brain slices were prepared as described previously (48). Briefly, following euthanasia, brains were removed and mounted onto a metal stage. Coronal slices containing mid-SCN levels across the rostro-caudal axis were cut using a Campden 7000smz-2 vibrating microtome (Campden Instruments, Loughborough, UK). Slices were cut at 250µm thickness for whole-cell recordings and at 350 µm for long term *in vitro* extracellular multi-electrode recording. Slicing was done in an ice-cold (4°C) sucrose-based incubation solution containing the following (in mM): 3 KCl, 1.25 NaH2PO4, 0.1 CaCl2, 5 MgSO4, 26 NaHCO3, 10 D-glucose, 189 sucrose, oxygenated with 95% O2, 5%CO2. After cutting, slices were left to recover at room temperature in a holding chamber with continuously gassed incubation solution for at least 20 min before transferring into aCSF. Recording aCSF has the following composition (mM): 124 NaCl, 3 KCl, 24 NaHCO3, 1.25 NaH2PO4, 1 MgSO4, 10 D-Glucose and 2 CaCl2, and 0 sucrose. For long term *in vitro* recordings, aCSF was supplemented with 0.001% gentamicin (Sigma-Aldrich). Slices were allowed to rest for at least 20 min before placement in the recoding chamber of the multielectrode array (MEA) or the upright microscope where they were continuously perfused with continuously gassed aCSF for the entire recording (∼3ml/min in the MEA and ∼2.5ml/min in the microscope).

### In vitro extracellular multi-electrode recordings

Recordings were performed using 60pMEA100/30iR-Ti-gr perforated multielectrode arrays comprising 59 electrodes (100 µm apart and arranged in a 6×10 layout, see Fig. 3A) as previously described (48). Optimal brain slice positioning and aligning onto the recording electrodes were confirmed by overlaying images captured with a GXCAM-1.3 camera (GX Optical, Haverhill, UK) fitted to a dissecting microscope. MEA recordings were performed at 34 °C.

Data were sampled at 50 kHz with MC_Rack Software using a USB-ME64 system and MEA 1060UP-BC amplifier from Multi-Channel Systems (MCS GmbH, Reutlingen, Germany). Data were then high-pass filtered at 200Hz and 2nd order Butterworth. Events/spikes were extracted by setting up a threshold (−17.5 µV) based on baseline noise level measured during tetrodotoxin (TTX) treatment. Baseline multiunit activity (MUA) levels were recorded for a period >24h. At the end of each experiment, the glutamate agonist NMDA (20uM) was bath applied for 5 min to confirm maintained cell responsiveness, followed by 1 μM TTX (∼8 min) to confirm acquired signals exclusively reflected Na^+^-dependent action potentials. All drugs were purchased from Tocris Bioscience (Bristol, UK) and stored as stock solutions (prepared in purified dH_2_O) at −20°C. Drugs were diluted to their respective final concentrations directly in pre-warmed, oxygenated aCSF on the experimental day.

### Whole-cell voltage- and current-clamp recordings

Coronal brain slices containing mid-level of the SCN (250 µm) were placed in the bath chamber of an upright Leica epi-fluorescence microscope (DMLFS; Leica Microsystems Ltd) equipped with infra-red video-enhanced differential interference contrast (IR/DIC) optics. Slices were held in placed with a tissue anchor grid, and continuously perfused with aCSF by gravity. Recordings were performed from neurons located across the whole SCN (see Fig. 4A) during the day and at night. SCN neurons were identified with a 40x water immersion UV objective (HCX APO; Leica) and targeted using a cooled Teledyne Photometrics camera (Retiga Electro). Photographs of the patch pipette sealed to SCN neurons were taken at 10X at the end of each recording for accurate confirmation of anatomical location of the recorded cell within the SCN.

Patch pipettes (resistance 7–10MΩ) were fashioned from thick-walled borosilicate glass capillaries (Harvard Apparatus) pulled using a two-stage micropipette puller (PB-10; Narishige). Recording pipettes were filled with an intracellular solution containing the following (in mM): 120 K-gluconate, 20 KCl, 2 MgCl2, 2 K2-ATP, 0.5 Na-GTP, 10 HEPES, and 0.5 EGTA, pH adjusted to 7.3 with KOH, measured osmolarity 295–300 mOsmol/kg). Cell membrane was ruptured at −70 mV under minimal holding negative pressure.

An Axopatch Multiclamp 700A amplifier (Molecular Devices) was used in voltage-clamp and current-clamp modes. Signals were sampled at 25 kHz and acquired in gap-free mode using pClamp 10.7 (Molecular Devices). Access resistance for the cells used for analysis was <30 MΩ and series resistance was below 50 MΩ. Post-synaptic currents (PSCs) were measured under voltage-clamp mode while holding the cells at −70mV. Measurement of spontaneous activity in current-clamp mode was performed with no holding current (I=0). All data acquisition and protocols were generated through a Digidata 1322A interface (Molecular Devices). All recordings were performed at room temperature (∼ 23°C).

### Data and statistical analysis

#### Analysis of behavioural and physiological data

General activity, WRA and Tb actograms were generated using El Temps (El Temps version 1.228; © Díez Noguera, University of Barcelona). Averaged mean waveforms and quantitative analysis of activity, sustained immobility and Tb patterns under LD conditions were calculated based on data (in 30 min bins) across the last 8 days of each lighting stage. Intradaily variability, a measure of rhythm fragmentation within a day, and day-to-day stability (interdaily-stability) for each rhythm were calculated as previously described (49). Sustained immobility (which in mice corresponds to episodes of sleep) was defined as a period of immobility > 40s, based on published criteria (45).

To calculate the phase angle of entrainment and circadian period of the activity rhythm, 3 experienced scorers, blind to the lighting conditions, fitted a line across activity onsets under constant dark conditions. Phase angle of entrainment was expressed as the predicted time of activity onset on the last day under LD conditions by extrapolation of the fitted line. Values reported in the manuscript are the average of those obtained by the three investigators. Percentage of variance (%V) in activity accounted for by the circadian period was determined by Sokolove-Bushell periodogram (El Temps version 1.228) and used as an indicator of rhythm robustness under DD. Relationship between daytime irradiance and different rhythm parameters was evaluated by linear regression analysis, with p<0.05 indicating that the slope of the fitted line is significantly non-zero.

#### Analysis of extracellular multi-electrode recordings

Long-term MEA recordings were analysed using custom Matlab routines as previously described (48). MUA recorded by each electrode/channel was considered to exhibited circadian variation when better fit by a sinusoidal function (constrained to a periodicity between 20 and 28h) than a first-order polynomial. Peak and trough firing for each rhythmic channel was determined from a 60s binned time-series (smoothed with a 2h boxcar filter). Percentages of rhythmic vs non-rhythmic channels between lighting conditions were compared using Fisher’s Exact Test.

#### Analysis of whole-cell recordings

PSCs frequency and amplitude (threshold of 5pA) analysis was performed offline by template-based sorting in Clampfit 10.7 (Molecular Devices) within a 30s window as previously described (50). Current-clamp data were analysed using Spike2 software (Cambridge Electronic Design, CED). Resting membrane potential (RMP), spontaneous firing rate (SFR) and input resistance (*R*_input_) were determined within 2 min of membrane rupture. Average SFR in firing cells was calculated as the number of action potentials per second within a 30s window of stable firing using a custom-written Spike2 script, and average RMP was measured as the mean voltage over a 30s window. *R*_input_ was estimated using Ohm’s law (R=V/I) where V represents the change in voltage induced by a hyperpolarizing current pulse (−30pA for 500 ms) as previously described (20). Percentages of cells in the different electrophysiological states during the day and at night from bright and dim conditions were analysed using Chi-Squared test.

#### Statistical analysis

Non-normal distributed electrophysiological data from different lighting conditions and time-of-day were compared using Mann-Whitney U Test. All statistical analysis were performed using SPSS version 23 (SPSS Inc., Chicago, IL, USA) and GraphPad Prism 7.04 (GraphPad Software Inc., CA, USA. For all tests, statistical significance was set at p<0.05. Data are expressed as mean ± SEM. Sample sizes are indicated throughout the text and figure legends.

## Acknowledgements

We would like to thank the members of the University of Manchester Biological Services Facility and Jonathan Wynne for their excellent assistance in colony maintenance and husbandry. We also thank Prof Simon Luckman for allowing us access to his electrophysiology equipment, and Dr Josh Mouland for his assistance with light measurements.

This work was funded by a Biotechnology and Biological Sciences Research Council (BBSRC) Industrial Partnership Award with Signify (BB/P009182/1) to RJL, and by grants from the BBSRC to TMB (B/N014901/1) and to MDCB (BB/S01764X/1), and the Wellcome Trust (210684/Z/18/Z) to RJL.

---

**Supplementary Table 1.** Behavioural and physiological rhythms under different daytime irradiances. Data are expressed as mean ± SEM (n=12); Significant p values (p<0.05) are in bold.

| n=12 | Log Mel Effective photon<br>flux/cm<sup>2</sup>/s | Dimmest<br>12.77 | Dim<br>12.95 | Mid<br>13.66 | Bright<br>14.80 | Is slope<br>significantly<br>non-zero? |
| --- | --- | --- | --- | --- | --- | --- |
| General<br>Activity | Mean Activity-Light | 390.40 ± 26.50 | 343.70 ± 16.08 | 392.50 ± 26.01 | 368.10 ± 20.39 | 0.997 |
|  | Mean Activity-Dark | 68.53 ± 7.14 | 69.21 ± 9.42 | 60.98 ± 7.26 | 73.73 ± 11.29 | 0.708 |
|  | Mean-24h | 229.50 ± 11.99 | 206.50 ± 8.68 | 226.80 ± 12.05 | 220.90 ± 11.21 | 0.885 |
|  | % Diurnality | 84.130 ± 2.614 | 83.300 ± 2.039 | 85.740 ± 2.202 | 83.350 ± 2.501 | 0.934 |
|  | Day-to day Reproducibility | 0.668 ± 0.037 | 0.638 ± 0.042 | 0.739 ± 0.031 | 0.718 ± 0.030 | 0.128 |
|  | Intradaily-Variability | 0.512 ± 0.057 | 0.542 ± 0.057 | 0.439 ± 0.057 | 0.446 ± 0.051 | 0.219 |
| Wheel<br>Running<br>Activity | Mean Activity-Light | 684.90 ± 71.64 | 598.60 ± 69.28 | 542.70 ± 77.12 | 625.60 ± 107.20 | 0.768 |
|  | Mean Activity-Dark | 39.13 ± 12.58 | 30.54 ± 12.31 | 33.94 ± 9.20 | 60.13 ± 13.97 | 0.113 |
|  | Mean-24h | 362.00 ± 36.60 | 314.50 ± 35.83 | 288.30 ± 39.67 | 342.90 ± 53.33 | 0.952 |
|  | Total Activity-Light | 16438.0 ± 1719.0 | 14365.0 ± 1663.0 | 13025.0 ± 1851.0 | 15015.0 ± 2573.0 | 0.768 |
|  | Total Activity-Dark | 939.2 ± 302.0 | 732.9 ± 295.4 | 814.6 ± 220.7 | 1443.0 ± 335.2 | 0.113 |
|  | Total Activity -24h | 17377.0 ± 1757.0 | 15098.0 ± 1720.0 | 13840.0 ± 1904.0 | 16458.0 ± 2560.0 | 0.952 |
|  | % Diurnality | 91.930 ± 3.675 | 94.920 ± 1.820 | 90.950 ± 4.215 | 87.130 ± 4.366 | 0.175 |
|  | Day-to day Reproducibility | 0.648 ± 0.039 | 0.621 ± 0.058 | 0.628 ± 0.036 | 0.646 ± 0.029 | 0.867 |
|  | Intradaily-Variability | 0.512 ± 0.103 | 0.466 ± 0.059 | 0.498 ± 0.059 | 0.534 ± 0.077 | 0.661 |
| Body<br>Temperature | Mean-Light | 37.98 ± 0.11 | 37.92 ± 0.09 | 38.12 ± 0.10 | 38.16 ± 0.11 | 0.090 |
|  | Mean-Dark | 35.62 ± 0.09 | 35.64 ± 0.11 | 35.57 ± 0.07 | 35.59 ± 0.08 | 0.725 |
|  | Mean-24h | 36.80 ± 0.08 | 36.78 ± 0.09 | 36.85 ± 0.07 | 36.88 ± 0.08 | 0.364 |
|  | Day-to day Reproducibility | 0.782 ± 0.025 | 0.781 ± 0.026 | 0.840 ± 0.016 | 0.844 ± 0.010 | **0.012** |
|  | Intradaily-Variability | 0.340 ± 0.027 | 0.346 ± 0.026 | 0.275 ± 0.024 | 0.259 ± 0.017 | **0.005** |
| Sustained<br>Immobility | Mean-Light | 0.264 ± 0.029 | 0.265 ± 0.018 | 0.212 ± 0.027 | 0.200 ± 0.016 | **0.020** |
|  | Mean-Dark | 0.768 ± 0.012 | 0.751 ± 0.022 | 0.771 ± 0.021 | 0.768 ± 0.027 | 0.768 |
|  | Mean-24h | 0.516 ± 0.011 | 0.508 ± 0.015 | 0.491 ± 0.016 | 0.484 ± 0.016 | 0.098 |
|  | Day-to day Reproducibility | 0.639 ± 0.035 | 0.620 ± 0.031 | 0.743 ± 0.029 | 0.745 ± 0.022 | **0.002** |
|  | Intradaily-Variability | 0.657 ± 0.056 | 0.692 ± 0.057 | 0.506 ± 0.061 | 0.508 ± 0.052 | **0.017** |

